# Seasonal dynamics of mixotrophic phytoplankton in a freshwater habitat revealed by single-cell sorting

**DOI:** 10.1101/2025.06.29.662170

**Authors:** Ying Wang, Qian Li

## Abstract

The nutritional strategy of phago-mixotrophy among eukaryotic phytoplankton has been widely recognized and characterized across various taxonomic lineages. While their ecological role as bacterivores in different aquatic environments is acknowledged, establishing a clear connection between mixotrophic activities and taxonomy within natural communities remains challenging. In this study, we utilized fluorescence-activated single-cell sorting techniques in combination with microscopic observation to investigate the diversity and seasonal variations of bacteria-feeding mixotrophs in a freshwater habitat. We identified four groups of mixotrophic phytoplankton using both methods: Pedinellales (Dictyochophyceae), *Cryptomonas* (Cryptophyceae), *Dinobryon* (Chrysophyceae), and a single-chloroplast bearing group within Chlorophyta. These groups accounted for 58-96% and 54-95% of total mixotrophs identified by the two methods, respectively. Seasonal variations demonstrated that Pedinellales and Chlorophyta were more abundant in spring and summer/autumn, while *Dinobryon* was only present in winter when temperatures were lowest and light intensity was moderate. Our findings, with six classes and twelve genera identified as bacterivorous mixotrophs, highlight the promising potential of single-cell techniques in aquatic plankton ecology. This approach reveals the speciation and feeding activities of phago-mixotrophs in situ, thereby enhancing our understanding of their functional ecological roles within microbial food webs.

## Introduction

Early studies of pelagic bacterivores, conducted before the 21st century, primarily focused on heterotrophs, including heterotrophic flagellates, ciliates, and some amoebas (Sanders et al., 2001, Calbet et al., 2001). As research progressed, it became evident that a broad suite of eukaryotic phytoplankton, referred to as photo-phago-mixotrophs (hereinafter referred to as ‘mixotrophs’), can also effectively remove bacteria through phagocytosis. These predation interactions between bacteria and bacterivorous mixotrophs serve as a vital link between prokaryotes and eukaryotes within microbial food webs. Investigations of the taxonomic composition, abundances, and grazing ecophysiology of these bacterivores will ultimately enhance our understanding of trophic transfer efficiency between bacterial and eukaryotic communities, food web dynamics, as well as carbon and nutrients cycling in various aquatic ecosystems.

Mixotrophs that have been characterized are mostly based on laboratory studies that used isolates, demonstrating oceanic mixotrophs are commonly associated with various groups from haptophytes (Jones et al., 1994; Legrand et al., 2001; Avrahami & Frada, 2020; Koppelle et al., 2022), dinoflagellates (Nygaard & Tobiesen, 1993; Jeong et al., 2005), chlorophytes (Maruyama & Kim, 2013; Bock et al., 2022; Pang et al., 2022), dictyochophytes (Li et al., 2021; 2022), and others. In comparison, mixotrophic bacterivores reported for freshwater environments have primarily focused on chrysophytes (e.g., *Ochromonas* sp., Sanders et al., 1990; Wilken et al., 2019, and *Dinobryon* sp., Caron et al., 1993; Princiotta et al., 2016) and cryptophytes (e.g., *Cryptomonas* sp., Costa et al., 2022).

Identifying bacterivorous mixotrophs directly in the field can be challenging for several reasons. First, mixotrophs are diverse both phylogenetically and functionally (Li et al., 2022; Edwards et al., 2023). Their nutritional strategies can vary not only between clades and genera but also on a species level. For instance, a study of *Micromonas pusilla* isolated from the Arctic Ocean suggested mixotrophy (McKie-Krisberg & Sanders, 2014), while another *Micromonas* species (*Micromonas Polaris*) displayed no evidence of feeding during laboratory studies (Jimenez et al., 2021). Second, research indicates that the grazing behaviors of mixotrophs are versatile and highly adaptable. These behaviors can change rapidly depending on environmental conditions such as nutrient availability, light, temperature and prey abundance (Wilken et al., 2019; Edwards et al., 2019; Blossom & Hansen, 2021). Consequently, the mere potential to be a mixotroph does not guarantee that active mixotrophic activities will occur in a given environment (Princiotta et al., 2023). For example, the freshwater bacterivorous group *Cryptomonas* has been identified as primarily autotrophic in a humic mesotrophic lake (Tranvik et al., 1989). However, they can also use phagotrophy effectively to supplement energy under the ice of Antarctic lakes during winter (Marshall & Laybourn-Parry, 2002).

Accurately distinguishing and identifying mixotrophs within natural communities often requires advanced molecular and enumeration techniques (Beinsner et al., 2019). Traditional methodology can be labor-intensive, generally involves microscopic observation and prey labeling techniques that utilize fluorescence (McManus & Fuhrman, 1986; Connell et al., 2020). Other approaches include the application of bromodeoxyuridine (Fay et al., 2013; Millette et al., 2021), and the study of rRNA genes through techniques such as catalyzed reporter deposition-fluorescence in situ hybridization (Unrein et al., 2015; Xiao et al., 2025) and RNA stable isotope probing (Frias-Lopez et al., 2009; Wilken et al., 2023). Recent research has applied fluorescence-activated flow cytometry enumeration techniques to retrieve mixotrophs on community level (Li et al., 2023), as well as sorting techniques designed for higher taxonomic level retrieval of mixotrophs in aquatic environments (Wilken et al., 2019; Florenza et al., 2024).

The investigation of *in situ* mixotrophs and their activities, despite facing certain challenges, has been highlighted as a key research priority in the field of aquatic mixotrophy ecology (Millette et al., 2023; Wilken & McManus, 2024). In this study, we aim to assess the taxonomic composition of active mixotrophs at higher taxonomic levels (e.g. species and genera) within a freshwater habitat. This will be achieved through a traditional method integrating grazing experiments with fluorescent prey surrogates and microscopic observation, as well as a novel technique using fluorescence-activated flow cytometry single-cell sorting and sequencing. We expect to observe distinct eukaryotic community compositions in both non-sorted (the whole community) and sorted water samples (active bacterivores), with seasonal variations present in both communities. By identifying the taxonomic composition of active bacterivores at the single-cell level, this study will demonstrate the application of single-cell techniques in aquatic ecology and will help address existing knowledge gaps regarding *in situ* mixotrophic activities in freshwater systems.

## Materials and Methods

### Study site and sample collection

Water samples were collected at four different seasons between 2023 and 2024 from a shallow, small mesotrophic lake located at 31.0**°** N and 121.4**°** E. We focus on bacteria grazers in this study and thus only small, nano-sized eukaryotes (< 20 µm) were included in most of our analysis. Approximately 2 L water was collected from the surface layer, pre-filtered with a 20 µm nylon sieve (to remove microzooplankton grazers and larger populations), and placed into clean transparent bottles. Spring, Winter, Summer, and Autumn samples were collected in April 2023, December 2023, July 2024, and October 2024, respectively. Sampling on each date was taken at approximately the same time, between 10 am and 12 pm to ensure comparable metabolic activities of eukaryotes.

Temperature, salinity, dissolved oxygen, and chlorophyll-*a* concentrations of the total phytoplankton community (including nano- and micro-sizes) were measured at the surface using sensors integrated on a vertical profiler (the YSI Pro DSS, Xylem Inc., USA). Photosynthetically available radiance levels were determined with a light quantum sensor of QSL-2100 (Biosperical, USA). Upon arrival at the lab (within one hour), approximately 50 mL water from the pre-filtered bulk sample was aliquoted into smaller bottles to measure dissolved inorganic nitrogen (including nitrate, nitrite, and ammonia) and phosphate concentrations using the Auto Discrete Chemical Analyzer (AQ 400; Seal Analytical, Germany). An additional 500 mL of pre-filtered samples were filtered through a 0.7 μm glass fiber filter (Whatman) to measure chlorophyll-*a* concentrations from the nano-sized phytoplankton community, using acetone extraction and turner fluorescence reading.

### Nano- and Pico-sized microbial abundances in situ

Approximately three 2 mL aliquots were fixed with glutaraldehyde at 0.5% final concentrations (Sigma-Aldrich), treated with 0.01% final concentration of Pluronic F68 to avoid cell attachments to the plastic tubes, flash frozen in liquid nitrogen, and stored with cryovials in -80**°**C. These vials were thawed at a later time, vortexed gently and run on a CytoFLEX flow cytometer (FCM) (Beckman Coulter, USA) for counting pico- and nano-sized pigmented eukaryotes (PE), non-pigmented heterotrophic flagellates (HF), and heterotrophic bacteria (Hbac). Because some HF are difficult to retrieve on FCM, they were stained with DAPI (Biotium) and enumerated on a membrane filter under epifluorescence microscopy (Olympus 61 X). The final results of HF abundances used in the study were from these microscopic observations. Similarly, PE was also observed under microscopy and their mean abundances averaged from both FCM and microscopy were used for further analysis.

### Eukaryotic community composition

An additional 1L of pre-filtered water sample was simultaneously filtered onto a 0.2 µm polycarbonate membrane filter (Whatman) to collect the natural microbiome DNA. Total environmental DNA was extracted using a column-based, DNeasy PowerSoil Pro Kit (47016) (Qiagen, Germany) and purified using VAHTS DNA Clean Beads (N411-03, Vazyme, China). Afterwards, the hypervariable V4 region of eukaryotic 18S rRNA genes (∼400 bp) was amplified with a universal pri mer pair of 547F (5’- CCAGCASCYGCGGTAATTCC-3’) and V4R (5’- ACTTTCGTTCTTGA TYRA-3’) (Stoeck et al., 2010). PCR products from each sample were purified and pooled in equal amounts, followed by pair-end 2 x 250 bp sequencing using the Illumina NovaSeq platform at Shanghai Personalbio Co., Ltd. Quality check and chimeras removal of raw amplicon sequence variants (ASVs) were conducted using DADA2 (Callahan et al. 2016). Filtered ASV sequences we re further classified with the SILVA database (Pruesse et al. 2007) using QIIME2 (Bolyen et al., 2019).

ASVs annotated as multi-cellular organisms (e.g., Vertebrata and Arthropoda) and parasites (e.g., Cryptomycota and Amoeba) that were not the focus of this study were excluded from further ana lysis. This filtration resulted in a final total sequence of 286,840 retrieved from four seasons belonging to eleven classified and one unclassified eukaryotic phyla (**Supplementary Table S1**). All partial 18S rRNA gene sequences retrieved from the total community were deposited in GenBank with BioProject number PRJNA1204479.

### Grazing experiments with single and multiple-prey types

In order to identify bacterivores from the *in situ* water samples, approximately 200 mL of pre-filtered water was amended with 1.0 µm carboxylate-modified polystyrene yellow-green fluorescent beads (catalog #L4655, Sigma Aldrich) at a final concentration of ∼2.0 ×10^5^ mL^-1^, accounting for 10% - 50% of the *in situ* bacterial concentrations. The 1.0 µm beads were chosen as the prey surrogates in our previous study because they retrieved the best fluorescence detection sensitivity on flow cytometry (Li et al., 2023). In this study, we also conducted multiple-prey experiments including 1.0 µm and 0.5 µm polystyrene yellow fluorescent beads (catalog #L5530, Sigma Aldrich), as well as live *Synechococcus* (strain WH7803) to assess possible feeding preferences or discriminations on the 1.0 µm beads.

Microscopic observations of prey preferences revealed differences among various bacterivores; however, all were able to effectively ingest the 1.0 µm beads (**Supplementary Fig. S1**). Consequently, the 1.0 µm beads were chosen as the standardized prey surrogates for all grazing experiments conducted in this study. This selection aimed to optimize fluorescence-activated cell sorting efficiency during flow cytometry analysis (Li et al., 2023). To minimize cell clumping and prevent surface attachments between cells and beads, Pluronic F-68 (Sigma Aldrich) was added to the water samples at a final concentration of 0.01% (Marie et al., 2014).

### Single cell sorting of active bacterivores

Several hours post the inoculation of a grazing experiment, unfixed and unstained sample aliquots (1∼2 mL aliquot) were run on FCM FACS Aria II (Beckman Dickinson, USA) to sort bacterivores that were actively feeding at the single-cell level. Bacterivores including mixotrophs and heterotrophs, were gated together based on high green fluorescence excited from the ingested beads (800-50,000 a.u. of FITC channel with ∼400 volta gain). A threshold between higher red fluorescence (>500 a.u. of PerCP-Cy5.5 channel with ∼300 voltage gain) and lower red fluorescence (100-500 a.u. of PerCP-Cy5.5 channel with ∼300 voltage gain) was further applied to increase separation between putative mixotrophs and heterotrophs.

Active bacterivores were targeted randomly across the entire gating area, and sorted into 96-well plates prefilled with 1µL PBS buffer, at a 10 µL/min sample flow rate. Approximately half of the 96-well plate (48∼56 wells) was sorted for each sampling season except for Summer 2024, totaling 160 sorted single cells. Upon finish of sorting, the plate was immediately sealed and stored in a -80 freezer till further processing.

### Single-cell genomic amplification

Sorted cells were thawed, following a lysis/denaturation and isothermal genomic amplification reaction using the DLB buffer of REPLI-g Single Cell Kit (Qiagen, catalog #150343). The kit uses REPLI-g sc Polymerase, an optimized formulation of the high-fidelity enzyme Phi 29 polymerase, to amplify complex genomic DNA using multiple displacement amplification (MDA) technology that provided highly uniform amplification across the entire genome. Though primarily used with mammal cells and bacteria, a few studies have successfully used it in marine protists (Chen et al., 2021; Wilken et al., 2019; Florenza et al., 2024). The isothermal strand-displacing reaction was performed following the product instruction. Specifically, the lysis and denaturation reaction (with 3 μL pre-prepared lysis buffer) was incubated at 65 °C for 10 min, then inactivated by adding 3 μL Stop Solution. The denaturant DNA material was afterward mixed with 29 μL Repli-g sc Reaction Buffer (contains primers, dNTP, and salt), 9 μL H_2_O and 2 μL REPLI-g sc Polymerase, incubated for 8 h at 30℃, followed with a stop step by decreasing the temperature to 65°C for 5 min.

### 18S rRNA genes phylogeny

Amplified genomic DNA (gDNA) was diluted at 100×, and used as the downstream PCR amplification template. The near-full length 18S rRNA gene of eukaryotes was amplified by PCR with the TaqMan polymerase PCR System (Mei5bio, MF848) using forward primer Euk63F 5′-ACGCTTGTCTCAAAGATTA-3 and Euk1818R 5′-ACGGAAACCTTGTTACGA-3′ (Lepere, 2011). Amplicons were purified and sequenced (Sanger) using the same forward PCR primer of Euk63F, yielding an overall gene sequence length of 840-900 bp.

For phylogenetic analyses, all sequences had >98% similarity were grouped into the same operational taxonomic units (OTUs). Homologous reference sequences were then retrieved from the NCBI GenBank based on BLAST similarity, with additional sequences from certain neighbor genera. Multiple sequence alignments including 22 OTUs, 18 homologous references, and ten addition al taxa were performed with MAFFT v7.450 using the G-INS-i algorithm in Geneious R11.1.5. Terminal sites that lacked data for any of the sequences were trimmed and any sites with greater th an 0.5 % gaps were removed from the alignment, which generated a final sequence length of 837 bases. Phylogenetic trees were constructed using MrBayes v3.2.6 in Geneious R11.1.5 (Huelsenbeck & Ronquist, 2001), implementing the GTR substitution model and the CAT approximation of rate heterogeneity, supported by two runs of four chains for 1,000,000 generations which were subsampled every 200 generations. The Bayesian majority consensus tree was further edited wit hin iTOL (Letunic & Bork, 2019).

All partial 18S rRNA gene sequences retrieved from cell sorting were deposited in GenBank wit h accession numbers PV348919-PV348940.

### Microscopic observation

One hour post each of the grazing experiment, 10-20 mL water aliquots were fixed with glutaraldehyde (0.5% final concentration), stained with DAPI (Biotium) and filtered onto a 2.0 µm pore size polycarbonate membrane filter (Whatman). Filters were mounted on glass slides and scanned under epifluorescence microscopy (Olympus 61X) with illumination settings for DAPI (Ex. 405 nm, Em. 400–480 nm), chlorophyll (Ex. 450 nm, Em. 575-IF), prey (Ex. 545 nm, Em. 605/55 nm), and a longpass filter BA575-IF for all fluorescence between green-yellow and red.

Nano-sized eukaryotic cells that contain one to six inherited, regular-shaped chloroplasts were categorized as pigmented eukaryotes, and those with ingested prey were counted as mixotrophs or mixotrophic bacterivores. Four populations at the genus or order level that harbor different numbers and shapes of chloroplasts were identified based on their unique morphology and characteristics, referring to various literature (e.g., *Dinobryon*, Stoecker et al., 2017; Jeong et al., 2023; *Cryptomonas*, Hoef-Emden & Melkonian, 2003; *Pedinellales,* Gerea et al., 2016; Koppelle et al., 2024; Florenza et al., 2024; Chlorophyta, Silva, 1999) and book chapters (Kristiansen & Andersen, 1986; Edward, 2008; Wehr et al., 2015; Jone et al., 2021).

These identified genera include *Dinobryon* belonging to Chrysophyceae (protoplasts housed in a cylindrical to funnel-shaped lorica with a stigma), Pedinellales belonging to Dictyochophyceae (three to six chloroplasts located at the periphery of cells), *Cryptomonas* belonging to Cryptophyceae (hexagonal periplast plates and phycobilin-emitted orange fluorescence), and parietal single-chloroplast bearing Chlorophyta with common chloroplast shapes resemble the moon, horseshoes, and cups etc. The remaining mixotrophic bacterivores were classified as “Other mixotrophs,” likely including mixotrophs from other species of Ochrophyta, Dinoflagellata, Chlorophyta, and Cryptophyceae. Inherited chloroplasts absent nano-sized flagellates were all classified as heterotrophs as species identification based on morphology is complex, and those with ingested prey were counted as heterotrophic bacterivores.

## Results

### Seasonality of physio-bio-chemical characteristics of the study region

The investigated lake is located at 31.0°N, 121.4°E, within a temperate climatological zone in eastern China (**Fig. 1a**). Hydrographic parameters of the investigated lake showed clear seasonal variations, with gradually increasing temperature from Winter (6.7℃) to Spring (14.8℃) and Autumn (21.0℃), and peaked in Summer (31.1℃). Overall higher underwater light availability (96.7-177.3 µmol m^-2^ s^-1^ PAR) was observed in Winter and Spring. The low underwater light intensity (17.3-34.3 µmol m^-2^ s^-1^ PAR) observed in Summer and Autumn was due to shading by large water plants (**Table 1**; **Fig. 1b**). Concentrations of nitrate and nitrite were high in Summer and Winter (19.6-39.5 µmol L^−1^) and low in Autumn (2.0-2.5 µmol L^−1^). Ammonia was overall low across seasons (<0.1 µmol L^−1^) except for Winter (13.3 µmol L^−1^) (**Table 1**).

**Fig. 1.**
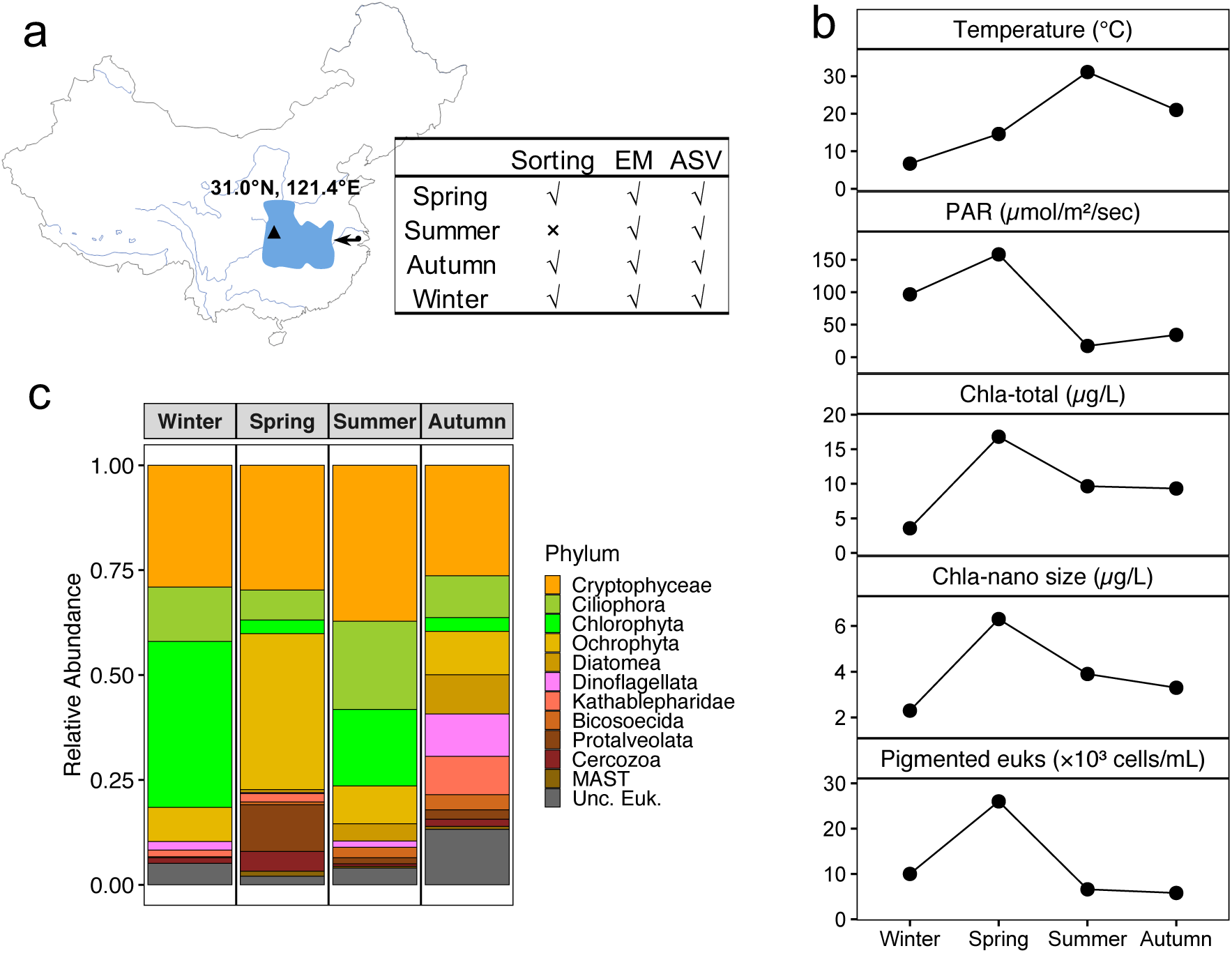
A map showing the geographic location of the study site, alongside a table listing the experimental design for each season (Sorting: single-cell sorting, EM: epifluorescence microscopy, ASV: amplicon sequencing) (a); plots presenting temperature, photosynthetically available radiation (PAR), chlorophyll a concentration of the total community, chlorophyll a concentration of the nano-sized eukaryotes, and abundance of nano-sized, pigmented eukaryotes across four seasons (b); phylum-level community compositions of total eukaryotes (including pigmented and aplastid) revealed by amplicon sequencing (c).

**Table 1.** Physical and biochemical characteristics of the study site.

| Season | PAR<br>($\mu\text{mol m}^{-2}\text{s}^{-1}$) | Temp<br>($^{\circ}\text{C}$) | $\text{NO}_3+\text{NO}_2$<br>($\mu\text{mol L}^{-1}$) | $\text{NH}_4$<br>($\mu\text{mol L}^{-1}$) | Chl *a*-Total<br>($\mu\text{g L}^{-1}$) | Chl *a*-Nano<br>($\mu\text{g L}^{-1}$) | PE<sup>a</sup><br>(cells $\text{mL}^{-1}$) | HF<sup>b</sup><br>(cells $\text{mL}^{-1}$) |
| --- | --- | --- | --- | --- | --- | --- | --- | --- |
| Winter | 96.7 | 6.7 | 19.6 | 13.3 | 3.6 | 2.3 | 1.0E+04 | 6.2E+03 |
| Spring | 158.2 | 14.6 | NA | NA | 16.8 | 6.3 | 2.6E+04 | 4.6E+03 |
| Summer | 17.3 | 31.1 | 39.5 | 0.1 | 9.7 | 3.9 | 6.6E+03 | 6.8E+03 |
| Autumn | 34.3 | 21.0 | 2.5 | 0.07 | 9.3 | 3.3 | 5.8E+03 | 6.1E+03 |

Abbreviations: Temp (temperature), PAR (photosynthetically available radiation), Chl *a*-Total (Chl *a* of total community), Chl *a*-Nano (Chl *a* of nano-sized eukaryote), PE (pigmented eukaryotes), and HF (heterotrophic flagellates).
<sup>a</sup>Mean value averaged between flow cytometry and microscopy-derived results
<sup>b</sup>Results were derived from microscopic observations only

The high light intensity and moderate temperature in Spring have introduced the highest chlorophyll-*a* concentration of both total community and nano-sized community (**Table 1**; **Fig. 1b**). Nano-sized pigmented eukaryotes (PE) also presented the highest abundance in Spring (2.6×10^4^ cells mL^−1^). Winter retrieved the lowest total Chl *a* concentration but the second highest PE abundance (1.0×10^4^ cells mL^−1^), possibly dominated by small pico-eukaryotes such as *Picochlorum* (**Supplementary Table S1**). Summer and Autumn retrieved similar Chl *a* and PE concentrations, and the lower PE concentrations could be partially due to light limitation. Heterotrophic flagellates (HF) demonstrated less variation across seasons, ranging between 4.6 × 10^3^ cells mL^−1^ in Spring, and up to 6.8 × 10^3^ cells mL^−1^ in Summer.

Total community composition revealed by amplicon sequencing suggested that pigmented eukaryotes were dominated by Chlorophyta, Cryptophyceae, Ochrophyta, Bacillariophyta, and Dinoflagellata. The majority of heterotrophs were composed of Ciliophora, Kathablepharidae, Bicosoecida, Cercozoa, MAST, and Protalveolata (**Fig. 1c**). Based on the 18S rRNA gene abundance, Cryptophyceae were abundant in all three seasons, always dominated by *Cryptomonas*. The second most abundant phylum was Chlorophyta, dominated by *Picochlorum* in Winter, and unclassified Chlorophyceae in Summer and Autumn (**Supplementary Table S1**).

### Single-cell sorting efficiency

The active bacterivore populations with high green fluorescence excited from ingested prey on FCM showed different shapes among different seasons. The most distinguishable and intact cloud of feeding bacterivores was observed from the Winter sample (**Fig. 2a**). The Spring and Autumn bacterivore populations presented similar distribution shapes (**Figs. 2b, c)**. With the BD FACS Aria II sorter, we sorted a total of 160 single cells from three different seasons (Winter, Spring, and Autumn) (**Fig. 2b**). Among all sorted samples, 140 cells were successfully amplified with a 10k bp genomic band, and 92 cells retrieved positive 18S rRNA genes (**Fig. 2d**).

**Fig. 2.**
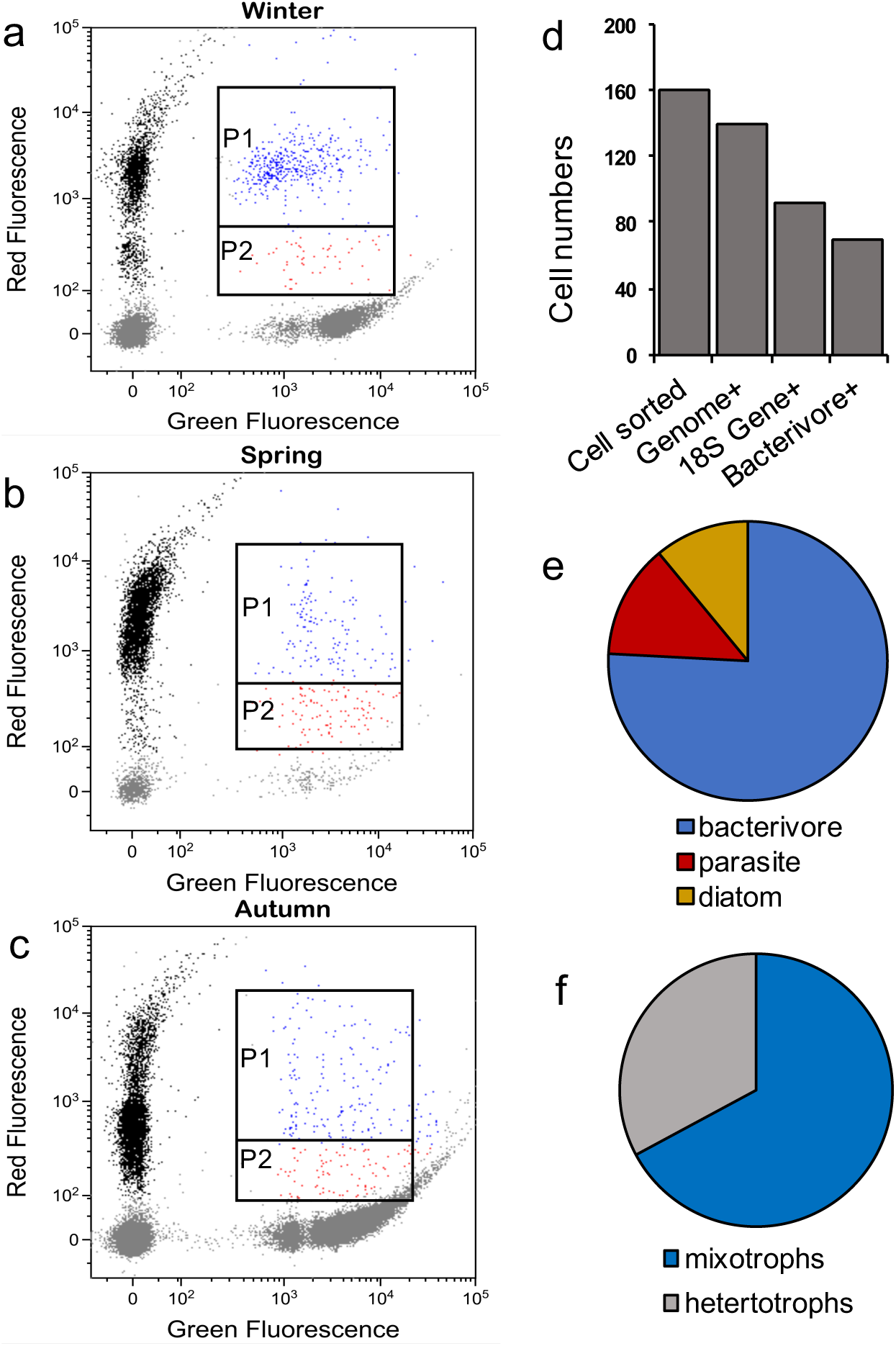
Flow cytometry graphs showing bacterivore populations gated for sorting putative mixotrophs (P1, events shown in blue) and putative heterotrophs (P2, events shown in red) with samples from Winter (a), Spring (b), and Autumn (c). Panel d demonstrates total events numbers from the initial sorting, samples recovered positive genomic amplifications, positive 18S rRNA genes, and annotated to be putative bacterivores. Panels e and f present community compositions in samples recovered with positive 18S rRNA genes and annotated to be bacterivores, respectively.

Out of those 92 eukaryotic cells, 70 cells were common bacterivores from aquatic environments, including 47 mixotrophs and 23 heterotrophs. The rest amplified 18S rRNA genes were affiliated with parasitic dinoflagellates and diatom, which were not considered bacterivores and excluded from further analysis (**Supplementary Table S2**; **Figs. 2e, f**). Consequently, our single-cell sorting retrieved a mean final success rate (number of identified bacterivores: total number of sorted single events) of 44% across all samples. Sorting efficiency varied significantly across seasons, possibly associated with community composition and bacterivorous activities. Specifically, Spring presented the highest final success rate of 73%, followed by Winter at 52% (19% false positive results from parasites), and Autumn at only 24% (26% false positive results from diatom).

Approximately 94% of all sorted mixotrophs were retrieved from the higher-chlorophyll P1 population on FCM and the other 6% were retrieved from the lower-chlorophyll P2 population on FCM (**Fig. 3**; **Supplementary Table S2**). In comparison, 61% of sorted heterotrophs were from the P2 population and the rest 39% were from the P1 population. Overall, the higher-chlorophyll P1 population revealed 83% mixotrophs, and the P2 population retrieved >90% heterotrophic bacterivores.

**Fig. 3.**
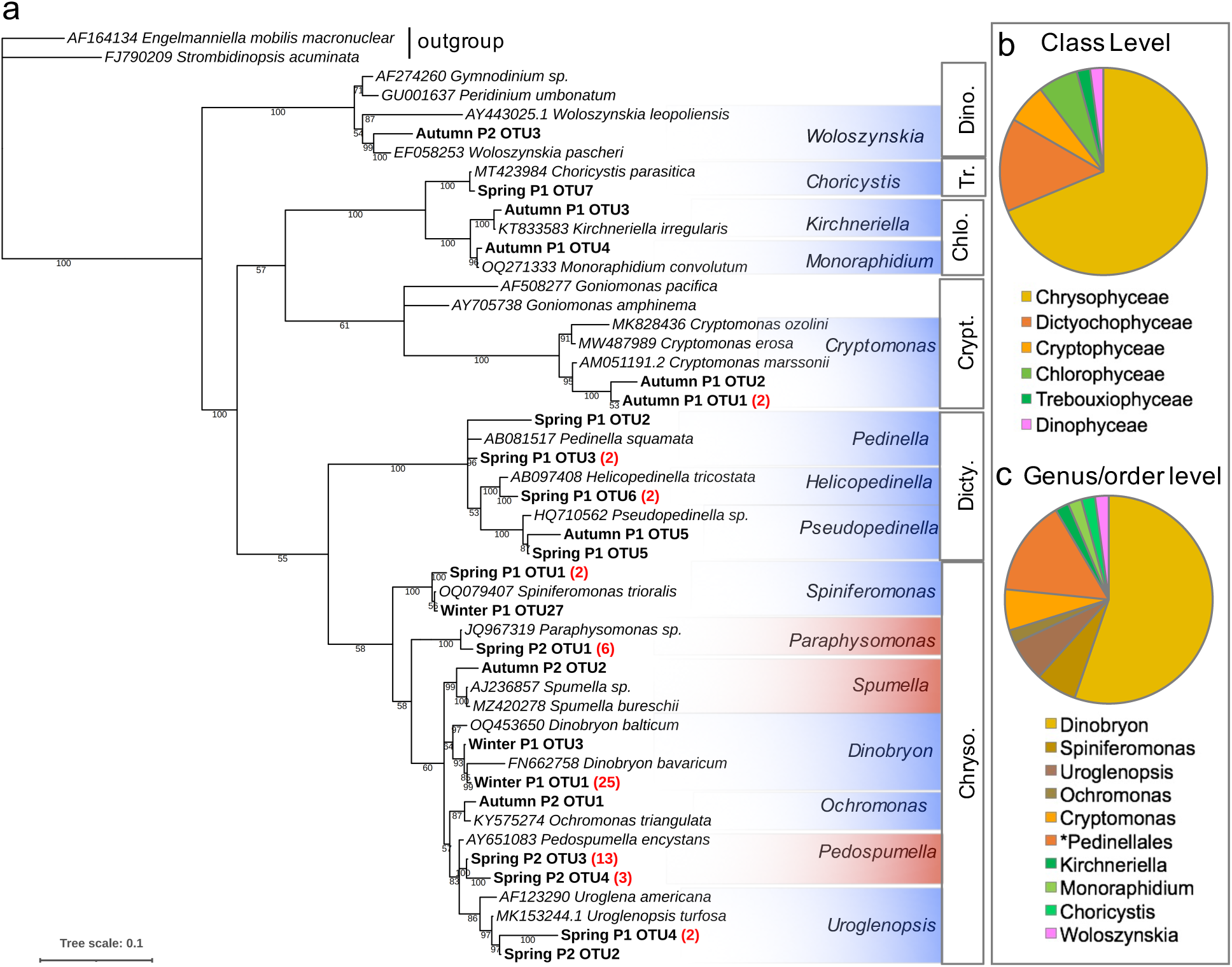
Neighbor-joining phylogenetic tree of sorted bacterivores constructed using partial 18S rRNA genes (a), and their taxonomic compositions on class level (b) and genus or order level (c) presented in pie charts. OTUs of bacterivores belonging to each genus in panel a were noted as Season_population gate on FCM_OTU#, followed by the total number of each OTU retrieved in a bracket if it was more than one. Abbreviations for class labels next to the tree in panel a were: Chryso. (Chrysophyceae), Dicty. (Dictyochophyceae), Crypt. (Cryptophyceae), Chlo. (Chlorophyceae), Tr. (Trebouxiophyceae), and Dino. (Dinophyceae). *Pedinellales was the only order-level group which included three genera of *Pedinella*, *Pseudopedinella*, and *Helicopedinella*. All OTU sequences are listed in Supplementary Table S2.

### Taxonomic composition of sorted bacterivores

These 70 successfully sorted and annotated bacterivores revealed 22 OTUs, belonged to 12 genera of mixotrophs and 3 genera of heterotrophs that were associated with six classes (**Fig. 3**). All 23 heterotrophic sequences (4 OTUs) were belonging to Chrysophyceae, composed of *Pedospumella*, *Spumella*, and *Paraphysomonas*. Focusing on only mixotrophs, Chrysophyceae accounted for 69% of total sorted bacterivores on class level (**Fig. 3b**). The second dominant class is Dictyochophyceae, contributing 15% of total sorted mixotrophic bacterivores. The rest four classes, Cryptophyceae, Chlorophyceae, Trebouxiophyceae, and Dinophyceae contributed 2-6% of total sorted bacterivores.

On genus or order level, the top two groups were *Dinobryon* of Chrysophyceae and Pedinellales of Dictyochophyceae, accounting for 55% and 15% of total bacterivores, respectively (**Fig. 3c**). The rest genera, including *Spiniferomonas, Uroglenopsis*, and *Ochromonas* of Chrysophyceae, *Pedinella*, *Pseudopedinella*, and *Helicopedinella* of Dictyochophyceae, *Monoraphidium* and *Kirchneriella* of Chlorophyceae, *Choricystis* of Trebouxiophyceae, *Cryptomonas* of Cryptophyceae, and *Woloszynskia* of Dinophyceae, contributed relatively less to all mixotrophic bactrivores, ranging from 2% to 6% of total community.

### Taxonomic composition of bacterivores observed through microscopy

Mixotrophic bacterivores were identified under epi-fluorescent microscopy from grazing experiments using single- and multiple-prey types, distinguished by their characteristic morphology and chloroplasts (see details in Methodology) (**Fig. 4a-g; Supplementary Fig. S1**). These mixotrophs included three groups at the genus/order level, including *Dinobryon* (Chrysophyceae), Pedinellales (Dictyochophyceae), and *Cryptomonas* (Cryptophyceae), and one group at the phylum level of Chlorophyta. Referring to the amplicon sequence and sorted bacterivore results, microscopy-identified single-chloroplast-bearing Chlorophyta mixotrophs were possibly associated with Chlorophyceae (such as *Kirchneriella* and *Monoraphidium*) and Trebouxiophyceae (such as *Choricystis*). The remaining mixotrophic bacterivores were classified as “Other mixotrophs”, possibly including mixotrophs from other populations of Ochrophyta (Chrysophyceae and Dictyochophyceae), Chlorophyta (Chlorophyceae, Trebouxiophyceae, and Mamiellophyceae), Dinophyceae, and Cryptophyceae.

**Fig. 4.**
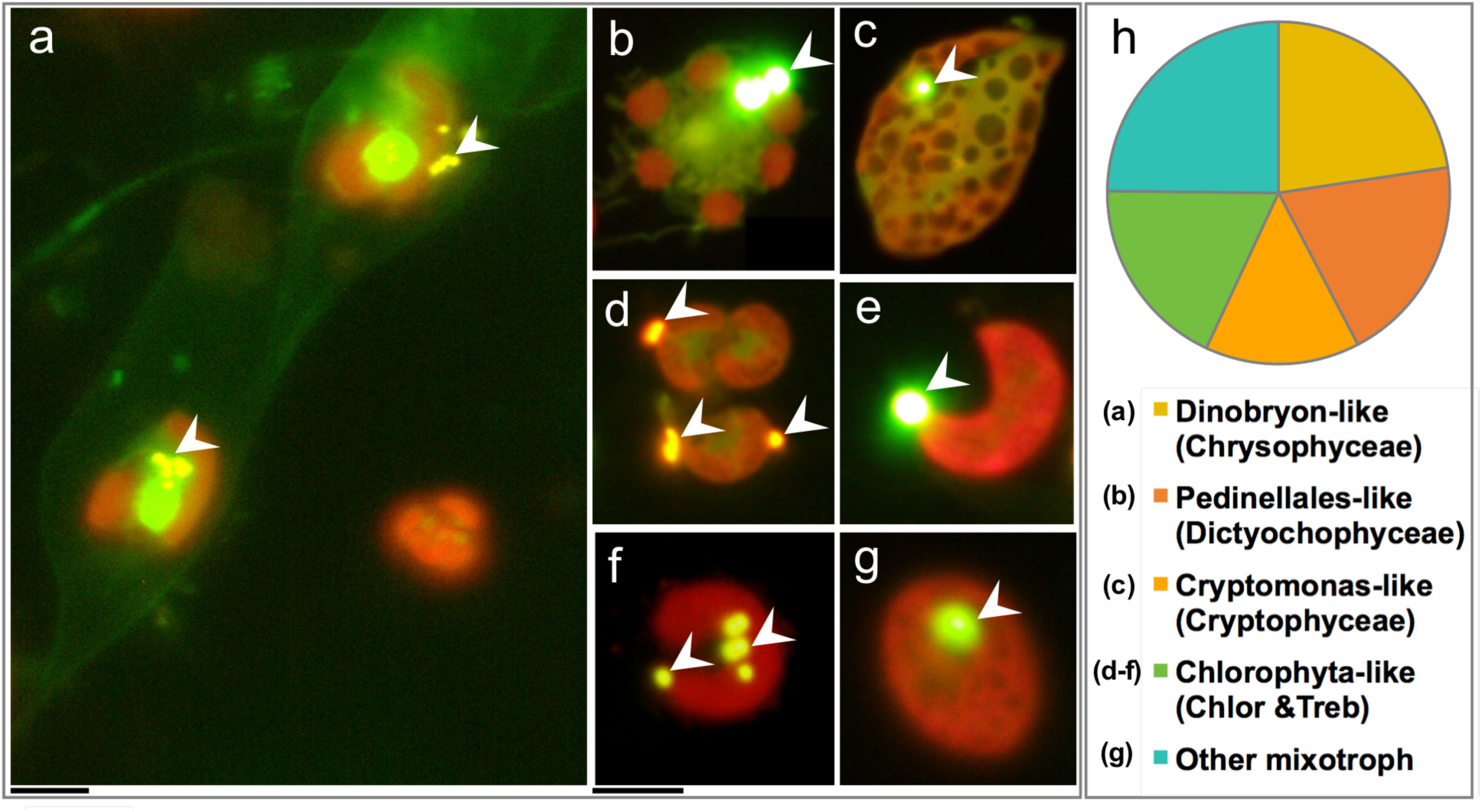
The five microscopy-identified mixotroph groups based on their distinguishable chloroplasts and morphology characters, including *Dinobryon*-like (a), Pedinellales-like (b), *Cryptomonas*-like (c), (single-parietal chloroplast-bearing) Chlorophyta-like (d-f), and “Other mixotrophs” did not have known unique morphological characteristics (g). Arrows denote ingestion evidence of various prey types (0.5 µm beads in a and f, *Synechococcus* in d, and all the rest were 1.0 µm beads). Proportions of each group across four seasons were shown in panel h. All scale bars are 5 μm in length; images b-g share the same scale bar beneath image f, and image a has its scale bar beneath image a. Legend of taxonomy in panel h were organized as genera/order (class); Chlor and Treb were abbreviations for Chlorophyceae and Trebouxiophyceae, respectively.

Among the five mixotrophic groups across different seasons, Dinobryon, “Other mixotrophs”, and Chlorophyta contributed 25%, 23%, and 18% of total bacterivores, which were moderately higher than the other three groups. Among Pedinellales and *Cryptomonas*, each group accounted for 20% and 15% of all observed bacterivores. Given that “Other mixotrophs” could encompass a much higher species diversity, those identified four mixotrophic groups were likely abundant, or common bacterivores across four seasons. All heterotrophic bacterivores were categorized as one single group of heterotrophs, and they contributed a similar proportion to total bacterivores across all seasons to mixotrophs (data not shown).

### Bacterivore compositions across different seasons obtained from two methods

All four identified bacterivore groups, observed microscopically, were also present in the sorted bacterivore communities, enabling a comparative analysis between the two approaches (**Fig. 5a, b**). Although the diversity of bacterivores retrieved through cell sorting was lower, partly due to the moderate to small sample size of sorted cells, the majority of common bacterivores were identified by both methods. This was particularly evident in the Autumn sample, where both methodologies identified the same four mixotrophic populations: (single-chloroplast bearing) Chlorophyta-like, *Cryptomonas*-like, Pedinellales-like, and “Other mixotrophs” (including other taxa from Chrysophyceae, Dinophyceae, Chlorophyceae, Trebouxiophyceae, Mamiellophyceae, Cryptophyceae, and Dictyochophyceae). In contrast, while Chlorophyta-like and *Cryptomonas*-like bacterivores were present in the microscopic samples collected during Winter and Spring, they were absent in the sorted community.

**Fig. 5.**
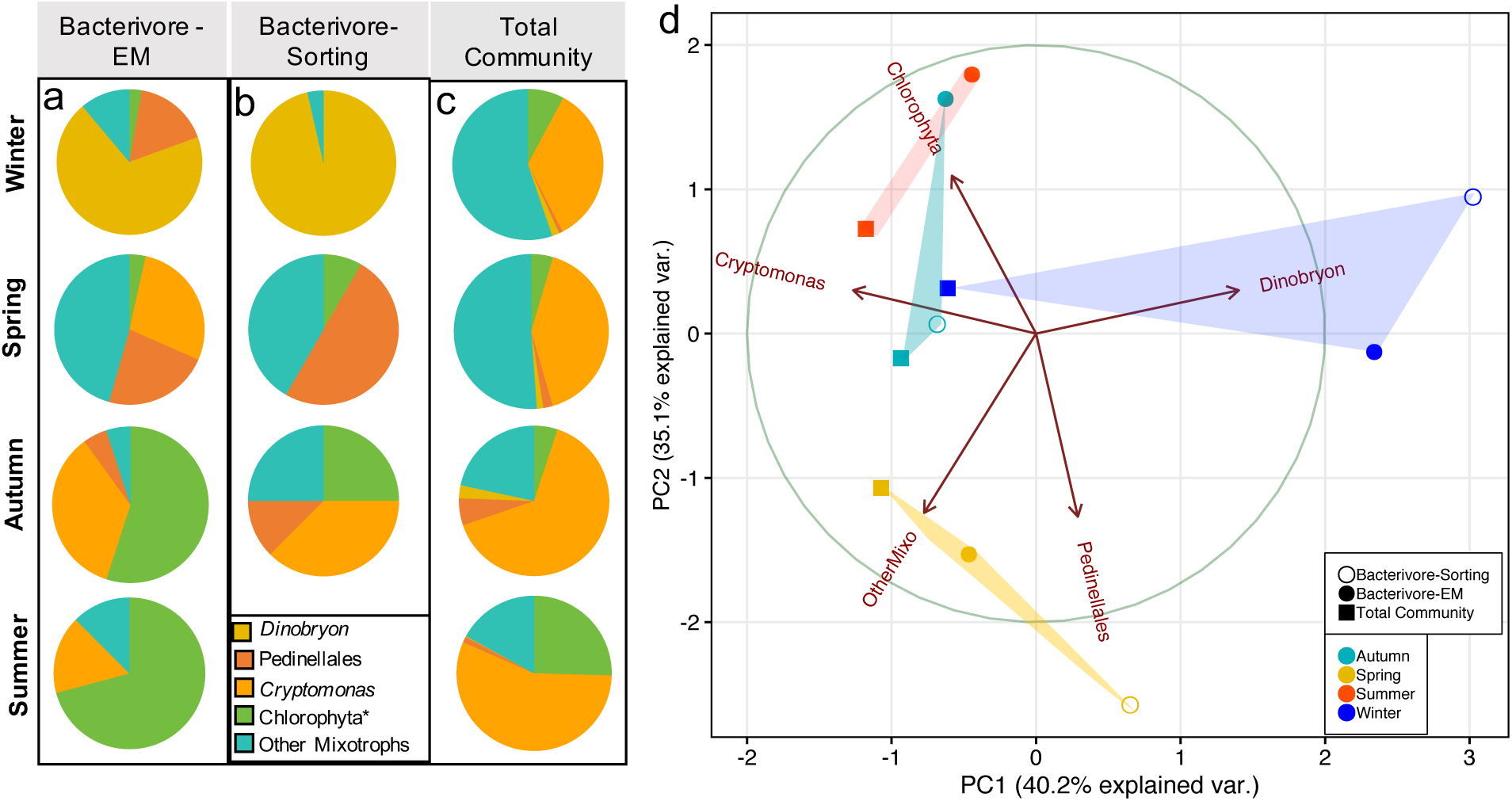
Pie charts demonstrating the taxonomic composition of five mixotrophic bacterivore groups obtained from epifluorescence-microscopy (EM) (a) and sorting approach (b) of each season. Panel c presents the relative abundances of those five groups in panels a and b in the total pigmented eukaryotic community (gene abundances of amplicon sequencing). Panel d was the principal component analysis plot showing Euclidean distances among all eleven samples in panels a-c. The circle in panel d denotes the PCA confidence ellipses, and samples from the same season were color-coded with shades. *A small portion of Chlorophyta taxa from the ASVs that likely contain more than one chloroplast were excluded from the analysis.

The bacterivore communities obtained through both methods demonstrated marked seasonal variations in composition, alongside significant differences when comparing to their relative abundances among the total pigmented-eukaryotic community (**Figs. 5a-c**). In the Winter season, *Dinobryon* emerged as the predominant bacterivore, detectable under microscopy and representing the sole group identified through flow cytometry. The Spring season featured notable presences of Pedinellales and “Other mixotrophs” in both sorted and microscopic analyses. During Autumn, both Chlorophyta and *Cryptomonas* were abundant across both methodologies. The Summer composition exhibited similarities to that of Autumn, with Chlorophyta identified as the most abundant bacterivores according to microscopic observations.

Principal component analysis (PCA) revealed similar patterns for the seasonality of bacterivore community composition (**Fig. 5d**). Specifically, the bacterivore community identified by microscopy in Summer and Autumn were positioned close to one another, exhibiting a strong correlation with Chlorophyta. Conversely, the Winter and Spring bacterivore communities were positioned further away, aligning more closely with *Dinobryon*, and Pedinellales and “Other mixotrophs”, respectively. Communities presented short distances with *Cryptomonas* encompassed sorted bacterivores in Autumn and total communities in Summer, Autumn, and Winter.

When heterotrophs were incorporated into the bacterivore community analysis, the Winter bacterivores were positioned away from the other three seasons, attributed to the much lower proportions of heterotrophic bacterivore in Winter (both microscopy and sorting results) (**Supplementary Fig. S2**). Distances between microscopy-identified bacterivores in Spring and Summer/Autumn were smaller than the PCA plot in Fig. 5, indicating similar proportions of heterotrophic bacterivores among these seasons. The positioning of different samples from the same season remained overall consistent with the PCA plot in Fig. 5. Nevertheless, sorted bacterivores in Spring presented a stronger association with heterotrophs and Pedinellales, while Summer/Autumn bacterivores identified through microscopy were positioned close to heterotrophs.

Lastly, when comparing the identified bacterivores to the total eukaryotic community across four seasons, the differences observed were more pronounced than the differences between the two methodologies used for bacterivore identification (**Figs. 5a-c**). These variations were consistent with the distances observed among samples based on principal component analysis, represented by the distances among all square symbols versus distances among all dots in **Fig. 5d**. Additionally, the distances between sorted bacterivore communities to the total community from the same season were overall larger than distances between two bacterivores identified by different methods, with the greatest distinction observed in Winter and the shortest in Autumn (between sorted bacterivore community and the total community).

## Discussion

### Common bacterivores in the investigated freshwater habitat

Identified mixotrophic bacterivores in the investigated freshwater habitat included members from the dictyochophytes, cryptophytes, chrysophytes, and chlorophytes. Among the three genera or orders identified through both cell sorting and microscopy, Pedinellales (dictyochophyte), *Cryptomonas* (cryptophyte), and *Dinobryon* (chrysophyte) all have been confirmed to possess phago-mixotrophic capabilities in various studies (Caron et al., 1993; Urabe et al., 2000; Gerea et al., 2016).

Kopelle et al. (2024) found that Pseudopedinella-like dictyochophytes and Cryptomonas-like cryptophytes were common bacterivores in a subalpine mountain lake. However, laboratory experiments have reported low ingestion rates in two species of *Cryptomonas* (Tranvik et al., 1989). Additionally, Gerea et al. (2016) demonstrated that Pseudopedinella were widespread across various lakes in higher latitudes of the Southern Hemisphere, contributing significantly to *in situ* bacterivory rates. In another temperate oligotrophic lake, Pedinellales also appeared to a dominant mixotrophic bacterivore group (Florenza et al., 2024).

For the single parietal plastid-bearing chlorophytes (*Kirchneriella* and *Monoraphidium*), the sample numbers in the sorted bacterivore community were low, with only one sequence/OTU identified for each genus. Therefore, any interpretations should be made with caution. Nevertheless, these chlorophytes were commonly identified as bacterivores through microscopy. Based on their cell morphology and plastid shapes, as well as (**Figs. 4d-4**), some of the ubiquitous bacterivores could possibly be *Kirchneriella* and *Monoraphidium* (the same as in sorted populations). Neither population was annotated in the overall community dataset, and it is possibly that some of the unclassified Chlorophyceae, which were dominant ASVs during summer and autumn, share similar morphological and physiological characteristics with these two mentioned genera.

Previous studies have demonstrated that various marine and freshwater species of chlorophytes possess mixotrophic capability, including members from Chloropicophyceae (Bock et al., 2021), Nephroselmidophyceae (Bock et al., 2021; Bell & Laybourn-Parry, 2003), and Pyramimonadophyceae (McKie-Krisberg & Sanders, 2014; Anderson et al., 2017), among others. However, none of the three genera of sorted Chlorophyta mixotrophs in this study, i.e., *Choricystis*, *Monoraphidium*, and *Kirchneriella*, have yet been confirmed to have phago-mixotrophic capabilities through laboratory studies.

### Seasonal variations of bacterivorous mixotrophs were observed in freshwater

Moderate to significant seasonal variations were observed for the different bacterivore populations identified via both cell sorting and microscopic approaches. Pedinellales, *Cryptomonas,* and chlorophytes were more commonly found in the identified bacterivore communities than *Dinobryon*, for which their seasonal variations were less likely caused by changes in relative abundances of each group. Rather, the observed seasonality could be caused by environmental condition shifts such as light, temperature, and nutrients, all of which could play an important role in determining the bacterivorous activities of different mixotrophic populations *in situ*.

For instance, the relative abundance of *Dinobryon* out of total eukaryotes (based on 18S rRNA gene copy numbers) was relatively consistent and low among seasons, ranging from 0.1% in Summer to 2% in Winter and 3% in Autumn (**Fig. 5c**). However, they were only identified as active bacterivore in Winter samples, accounting for 69% of microscopy observed bacterivores and 100% sorted bacterivores. This marked increase of dominance as bacterivores or mixotrophic activities of *Dinobryon* in Winter was associated with the lowest temperature (∼7℃) and second highest irradiance (∼100 µmol m^-2^ s^-1^) compared to other seasons throughout the year (**Table 1**). Using the culture of *Dinobryon sociale*, experimental studies have also suggested that bacterivory/mixotrophy of *Dinobryon* benefited from relatively low temperature (higher ingestion rates under 8-16℃ than 20℃) and relatively high irradiance (higher ingestion rates under 130 µmol m^-2^ s^-1^ than 66 µmol m^-2^ s^-1^) (Princiotta et al., 2016). Based on these results, *Dinobryon* seemingly benefits from bacterivory in colder seasons and under sufficient light for additional nutrients and energy supply (Kamjunke et al., 2006; Heinze et al., 2013).

Despite being overall ubiquitous bacterivores, abundances of Pedinellales and *Cryptomonas* demonstrated somewhat seasonality. Pedinellas were abundant or dominant in both microscopy- and sorting-derived communities in Spring, and *Cryptomonas* were abundant or dominant in Autumn. The increased Pedinellales bacterivory in Spring (higher temperature, ample irradiance, and high chlorophyll concentrations) agrees with previous findings suggesting that increased prevalence of Pedinellales was associated with high concentrations of total chlorophyll (Kopelle et al., 2024). In comparison, bacterivory/mixotrophy of *Cryptomonas* seemed to be affected by different environmental variables. For example, enhanced abundances of cryptophytes have been found to be associated with low light conditions (Teleaulex sp.; Jeong et al., 2005) and *in situ* mixotrophic activities (Tranvik et al., 1989). These observations agree with our results that *Cryptomonas* seemed to obtain a higher fitness when the light was low or limiting in Autumn and Summer (**Table 1**).

The last marked seasonal differences were found for the single parietal plastid-bearing mixotrophic chlorophytes, presenting a significantly higher proportion in the Summer and Autumn bacterivore communities, especially Summer. As mentioned above, one of the most distinct environmental conditions observed for those two seasons was the low underwater irradiance. However, physiological experiments under changing light conditions performed with chlorophyte species such as *Picochlorum* (Pang et al., 2023) and *Micromonas* (McKie-Krisberg & Sanders, 2014) have suggested that bacterivory were overall enhanced when they received ample light. This discrepancy could be explained by different grazing physiology of different chlorophyte species, or other variables *in situ* within a complex natural environmental setting, such as decreased nutrient concentrations in Autumn, and higher temperature observed in both seasons.

### Strengths and shortcomings of the single-cell sorting methodology

The average success rate of retrieved bacterivores (out of total sorted events) was approximately 50% across different seasons, among which around 60% were mixotrophs and 40% were heterotrophs. Using the same fluorescence-activated cell sorting technique, Florenza et al. (2024) have retrieved a larger number of mixotrophic bacterivores (943 bacterivores out of 1837 sorted single events) with higher diversity (34 OTUs across six classes) in a temperate oligotrophic lake. Nevertheless over 80% of their sorted bacterivores were Pedinellales, presented a 51% success rate that were comparable to this study. Collectively, the fluorescence-activated single cell sorting approach appeared to be highly efficient to retrieve the dominant bacterivores, showcased both successes and inherent limitations in uncovering mixotrophic diversity from natural aquatic environments.

The observed 50% false positive rate was primarily attributed to sorting failures, issues with cell lysis, and contamination during single-cell genomic amplification, each factor contributing to a 6-12% false positive rate. In addition, unintended interactions or attachment with beads other than phagocytosis due to cell surface stickiness (Seymour et al. 2017; Wilken et al., 2019) may explain for the sorted diatom and dinoflagellate parasitoid of *Pararosarium* (**Supplementary Table S2**), neither of which were seemingly to be bacterivores. It is speculated that the model of the flow cytometry sorter might influence sorting efficiency, and enhancements in cell lysis techniques could boost genomic amplification rates, particularly for groups such as chlorophytes with cellulose cell walls and dinoflagellates with thecas. Additionally, minimizing cell attachment could potentially be achieved by using non-ionic surfactants (Marie et al., 2014), reducing incubation time, and incorporating microscopy checks (Wilken et al., 2019).

Lastly, our findings indicate that heterotrophs that may also exhibit omnivorous behavior could be detected with positive chlorophyll fluorescence, such as *Spumella*-like flagellates (Silva, 1960). When focused on mixotrophs, specificity might be improved by adjusting and refining the gating area for red fluorescence values. Our results further suggested that sorted diatoms are associated with high chlorophyll fluorescence, while aplastid bacterivores often display lower chlorophyll fluorescence.

Collectively, this study applied fluorescence-activated single-cell sorting technique, combined with single-cell genomic amplification and microscopic observation into aquatic mixotrophy investigations, showcasing the potential for uncovering the broad diversity of pahgo-mixotrophs in natural water samples on single-cell levels. Our findings shed light on the ubiquity of phago-mixotrophic activities of pigmented eukaryotes in freshwater habitats, providing insights into their functional role mediating elements cycling between prokaryotes and eukaryotes within the aquatic food web.

## Data Availability Statement

All genetic sequence data obtained in this study were deposited in the NCBI GenBank (BioProject No. PRJNA1204479 and Gene Accession No. PV348919-PV348 940) (https://www.ncbi.nlm.nih.gov/), and can be obtained from the Supplementary Material. En vironmental variables and experimental results can be obtained from the main text, or upon reque st made to the authors.

## Acknowledgments

We acknowledge the funding agent for this study of the National Natural Science Foundation of China (Grants No. 42476120; 42106097), the UN Global Ocean Negative Carbon Emissions (ONCE) program, and the Ministry of Science and Technology (MOST) ONCE project. We also thank Dr. Haixia Jiang for operating the flow cytometry and the Research Facility Sharing Platform under the School of Life Sciences and Biotechnology at Shanghai Jiao Tong University.

## Conflicts of Interest

The authors declare no conflicts of interest.

## CRediT

Conceptualization - Q.L.; Data curation - Y.W. and Q.L.; Formal analysis - Q.L. and Y.W.; Funding acquisition - Qian Li; Investigation - Y.W. and Q.L.; Methodology - Q. L. and Y.W.; Supervision - Q.L.; Validation - Q.L.; Writing - original draft - Y.W. and Q.L.; Writing - review & editing - Q.L.

## Notes

### Competing Interest Statement

The authors have declared no competing interest.

