## Supplementary Material for "Seasonal dynamics of mixotrophic phytoplankton in a freshwater habitat revealed by single-cell sorting"

Supplementary material for “Seasonal dynamics of mixotrophic bacterivores in a freshwater habitat revealed by single-cell sorting” by Ying Wang and Qian Li

**Table legend** (in separate files)
Table S1. Amplicon sequence variants (ASVs) of partial 18S rRNA genes from total eukaryotic communities across four seasons. Datasheets were labeled as ‘season_original data’ and ‘season_filtered data’ which refer to sequences before and after removing non-targeting taxa such as parasites and metazoan.

Table S2. Taxonomic composition of all sorted events, including putative bacterivores and other protists (blue color-coded). Sequences from the same OTUs were only shown once. Sequence identifiers were noted as Season_population gate_OTU#.

**Figure legend**
Fig. S1. Comparison of ingestions by different mixotrophic groups among three prey types, including live *Synechococcus*, 1.0μm latex beads, and 0.5μm latex beads (a). Results were retrieved and averaged from multi-prey experiments conducted in different seasons. Observed prey numbers ingested by each mixotroph group were normalized against numbers of ingested 1.0μm beads. Mixotrophic groups were consistent with those shown in Fig. 4. The ‘x’ symbol notes the absence of ingestion of *Synechococcus* by *Dinobryon* from observed samples. Panel b shows ingestion evidence from representative cells from each mixotrophic group on different prey types.

Fig. S2. Pie charts demonstrating the taxonomic composition of all bacterivore groups identified from epifluorescence-microscopy (EM) (a) and cell-sorting (b) of each season, including five mixotrophic groups and one heterotrophic group. Panel c presents the relative abundances of those six groups in panels a and b in the total eukaryotic community (gene abundances of amplicon sequencing). Panel d was the principal component analysis plot showing Euclidean distances among all eleven samples in panels a-c. The circle in panel d denotes the PCA confidence ellipses, and samples from the same season were color-coded with shades. *A small portion of Chlorophyta taxa from the ASVs that likely contain more than one chloroplast were excluded from the analysis.

Fig. S1

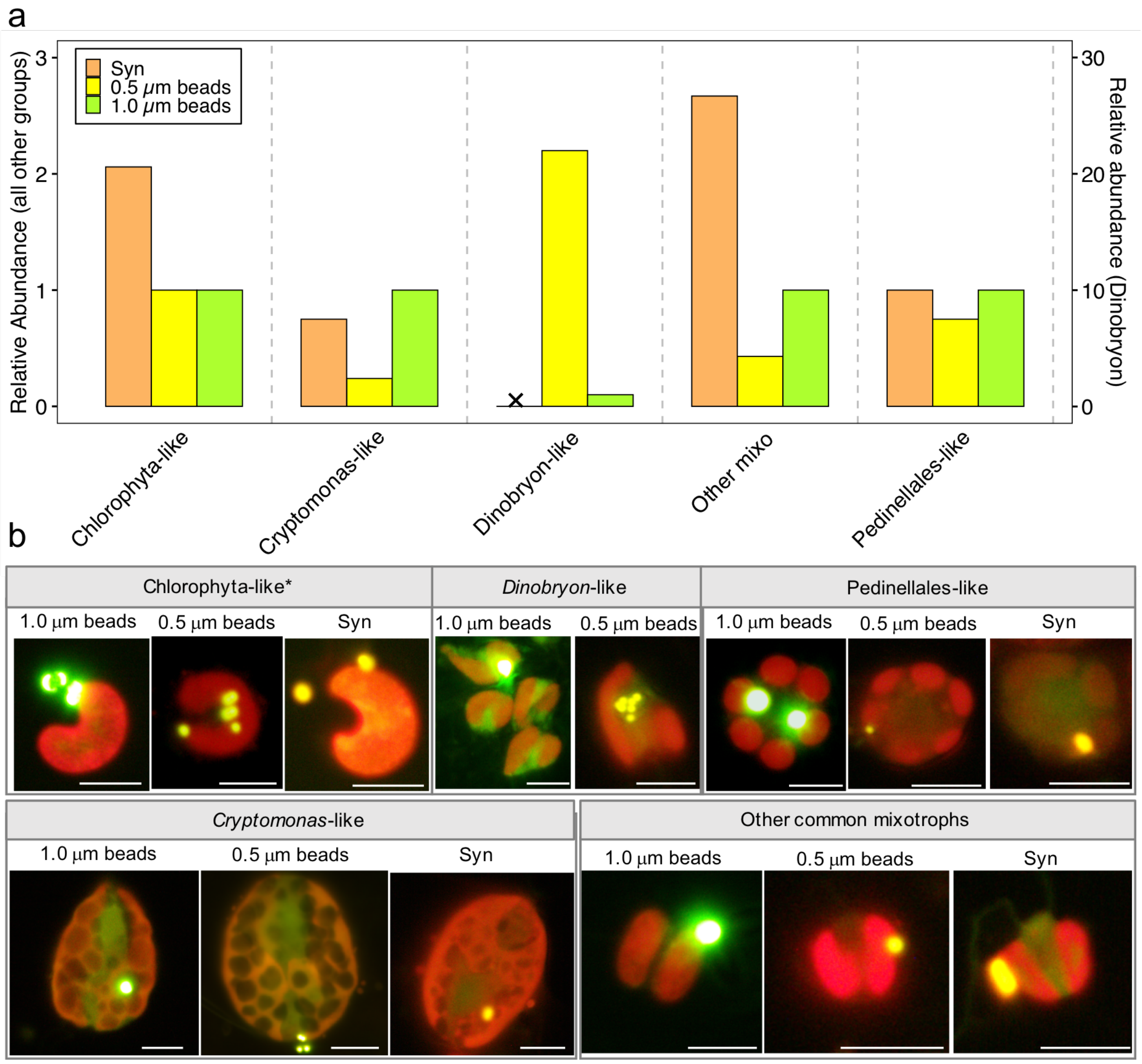
a
Syn
0.5 μm beads
1.0 μm beads
Relative Abundance (all other groups)
Relative abundance (Dinobryon)
0
1
2
3
0
10
20
30
Chlorophyta-like
Cryptomonas-like
Dinobryon-like
Other mixo
Pedinellales-like
b
Chlorophyta-like*
Dinobryon-like
Pedinellales-like
1.0 μm beads
0.5 μm beads
Syn
1.0 μm beads
0.5 μm beads
1.0 μm beads
0.5 μm beads
Syn
Cryptomonas-like
Other common mixotrophs
1.0 μm beads
0.5 μm beads
Syn
1.0 μm beads
0.5 μm beads
Syn

Fig. S2

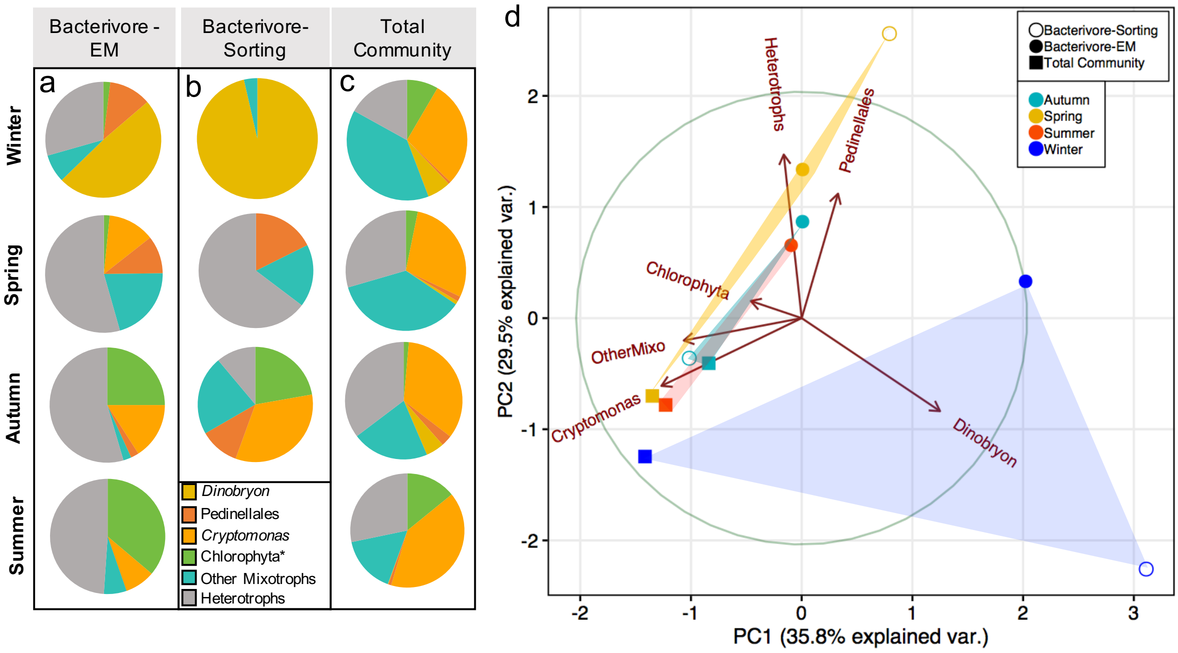

Bacterivore - EM
Bacterivore- Sorting
Total Community
a
b
c
d
Winter
Spring
Autumn
Summer
Dinobryon
Pedinellales
Cryptomonas
Chlorophyta*
Other Mixotrophs
Heterotrophs
Bacterivore-Sorting
Bacterivore-EM
Total Community
Autumn
Spring
Summer
Winter
Heterotrophs
Pedinellales
Chlorophyta
OtherMixo
Cryptomonas
Dinobryon
PC1 (35.8% explained var.)
PC2 (29.5% explained var.)
